## Supplementary Information for "Revealing intact neuronal circuitry in centimeter-sized formalin-fixed paraffin-embedded brain"

### Supplementary Tables

**Table S1**: The antibodies used in the study.
**Table S2**: Abbreviations of brain regions mentioned in Fig. 2B.
**Table S3**: The antibody conditions in the multi-round immunolabeling shown in Fig. 4.

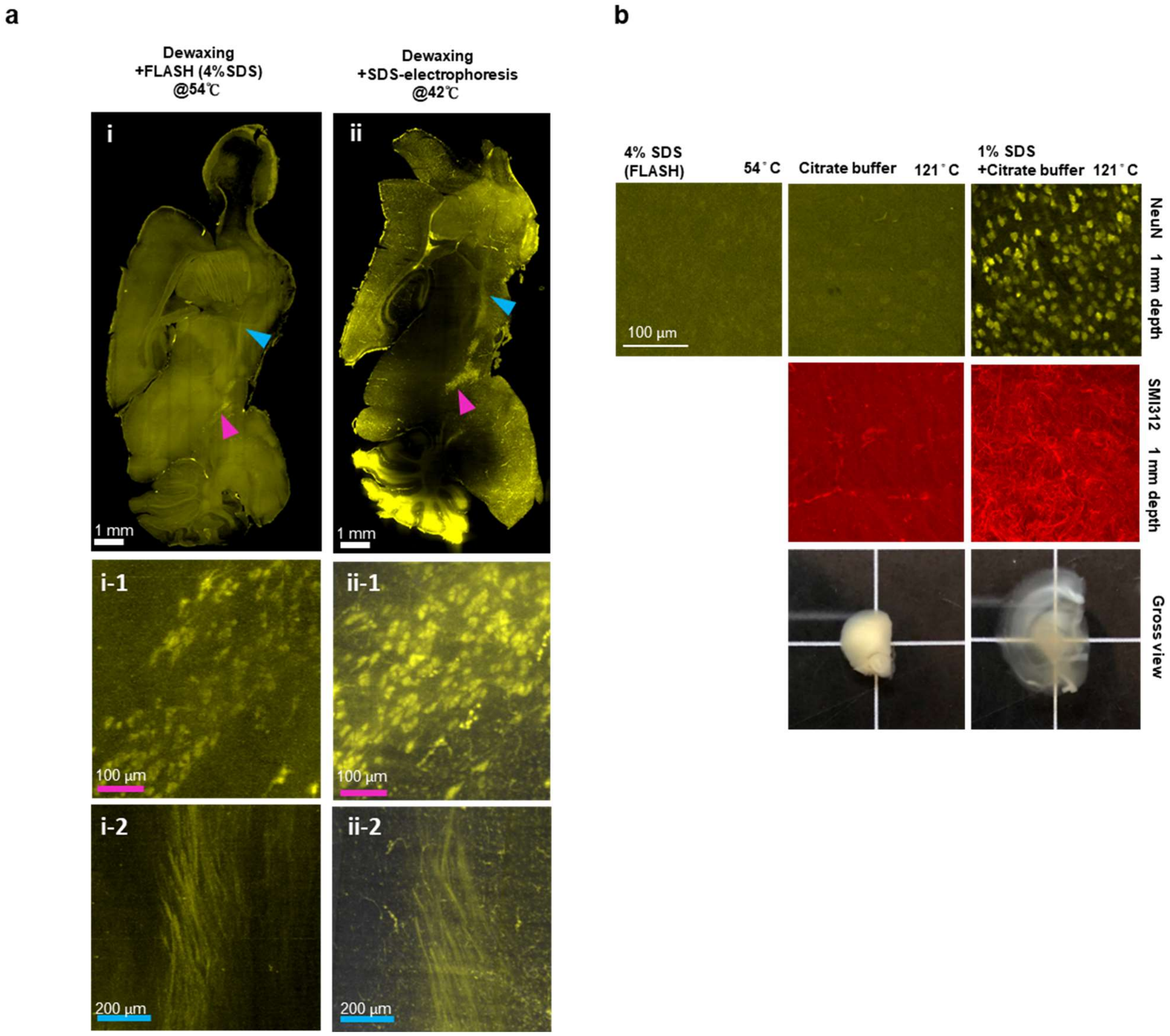


**Figure S1: Examination of the antigen retrieval effect of SDS on FFPE specimens.**

**a,** Light-sheet images of FFPE mouse brain hemisphere stained with Tyrosine hydroxylase (TH) antibodies. The SDS treatment conditions are described above the images. Magnifications of dopaminergic cells (magenta arrowheads) and nigrostriatal fiber tracts (cyan arrowheads) indicated in **i** and **ii** are shown in panels **i/ii-1** and **i/ii-2**, respectively.

**b,** Multi-point confocal fluorescence images of neuronal nuclei in the cortex (top row) and axons in the striatum (middle row), and gross view (bottom row) of 2-mm-thick FFPE mouse brain slices treated under various SDS conditions. Images of the optical section at a depth of 1 mm are shown. NeuN, a neuronal nuclear marker; SMI312, a pan-axonal marker.

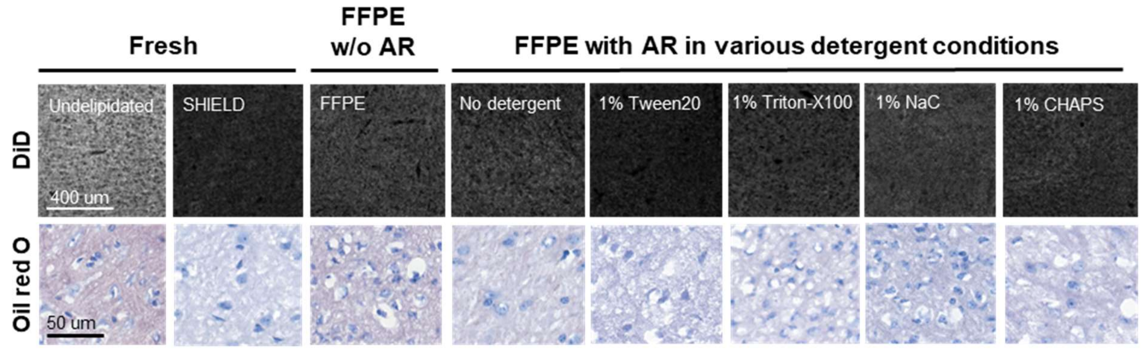


**Figure S2: Representative images of the results of DiD and Oil red O staining.** The delipidation effects of various detergent conditions were visualized and evaluated by lipophilic dye staining. Representative images of DiD (top row) and Oil red O (bottom row) staining are shown. A ZEISS 780 confocal microscope was used for DiD staining, whereas a 3DHISTECH panoramic slide scanner was employed for Oil red O staining.

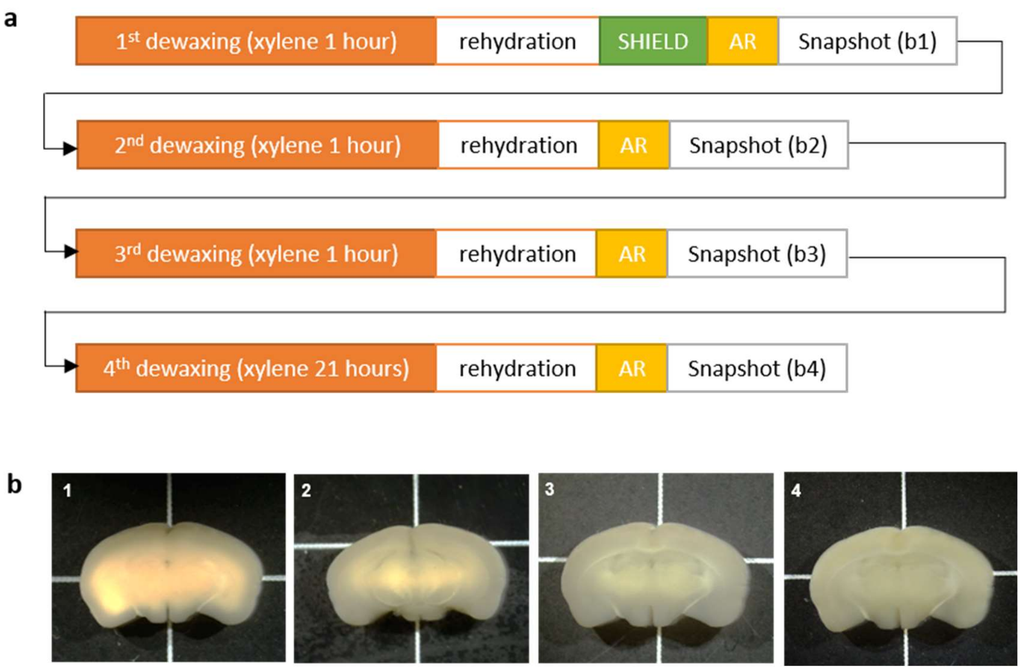


**Figure S3: Repeated Dewaxing Test.** This test was designed to mimic experimental conditions where the sample is large and the optimal dewaxing time needs refinement.

**a,** Test procedure: A 2-mm-thick mouse brain block was fixed with 4% PFA for 24 hours and then embedded in paraffin wax. The FFPE brain block was incubated at 65 °C to melt the paraffin and then immersed in xylene for 1 hour ("1st dewaxing"). The dewaxed brain block was rehydrated, SHIELD-processed, and subjected to antigen retrieval as described under the *HIF-Clear pipeline* subheading in the Materials and Methods. Consequently, a sufficiently dewaxed region turned translucent (b-1, the extremities of the brain), whereas any region with residual wax remained opaque (b-1, the central area of the brain). The insufficiently dewaxed brain block was then dehydrated and immersed in xylene for another hour ("2nd dewaxing"). The now twice-dewaxed sample was rehydrated, again subjected to antigen retrieval, and photographed. This process was repeated and, at the end of each cycle, a gross view image was acquired.

**b,** Gross views of the brain block after each dewaxing step. Note that the extent of the translucent border area increases while that of the opaque central area decreases with repeated dewaxings.

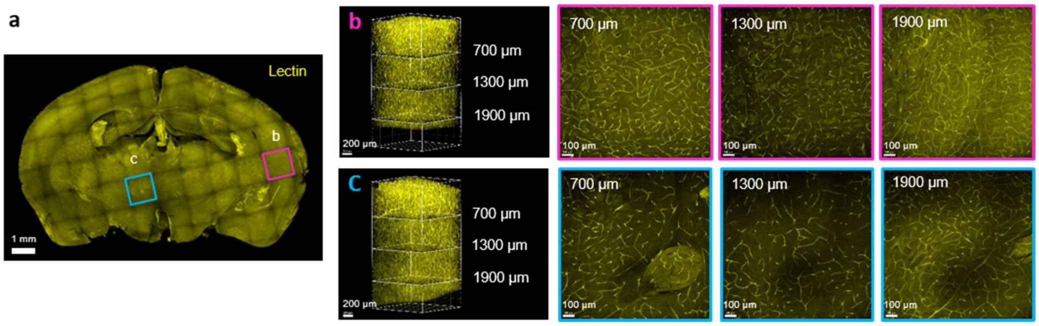


**Figure S4: Repeated dewaxing does not influence antigenicity.** The brain block subjected to four rounds of dewaxing (as shown in Supplementary Figs. 3b, 4a) was labeled with lectin to examine antigenicity.

**a**, Projection image of the entire block.

**b**, 3D image and optical sections of the magenta-framed box in **a**.

**c**, 3D image and optical sections of the cyan-framed box in **a**.

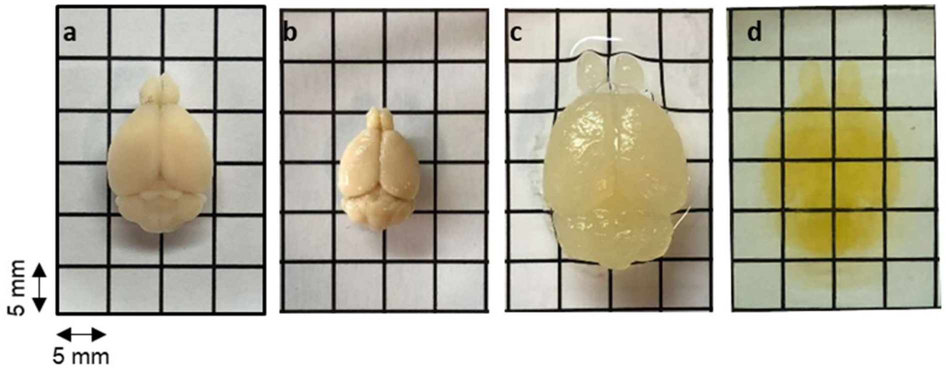


**Figure S5: Gross view of the mouse brain at different stages of the HIF-Clear protocol.**

**a,** 4% PFA fixation.

**b,** FFPE embedding.

**c,** optimized antigen retrieval.

**d,** RI-matching.

The FFPE mouse brain remain intact and turn transparent after processing with HIF-Clear.

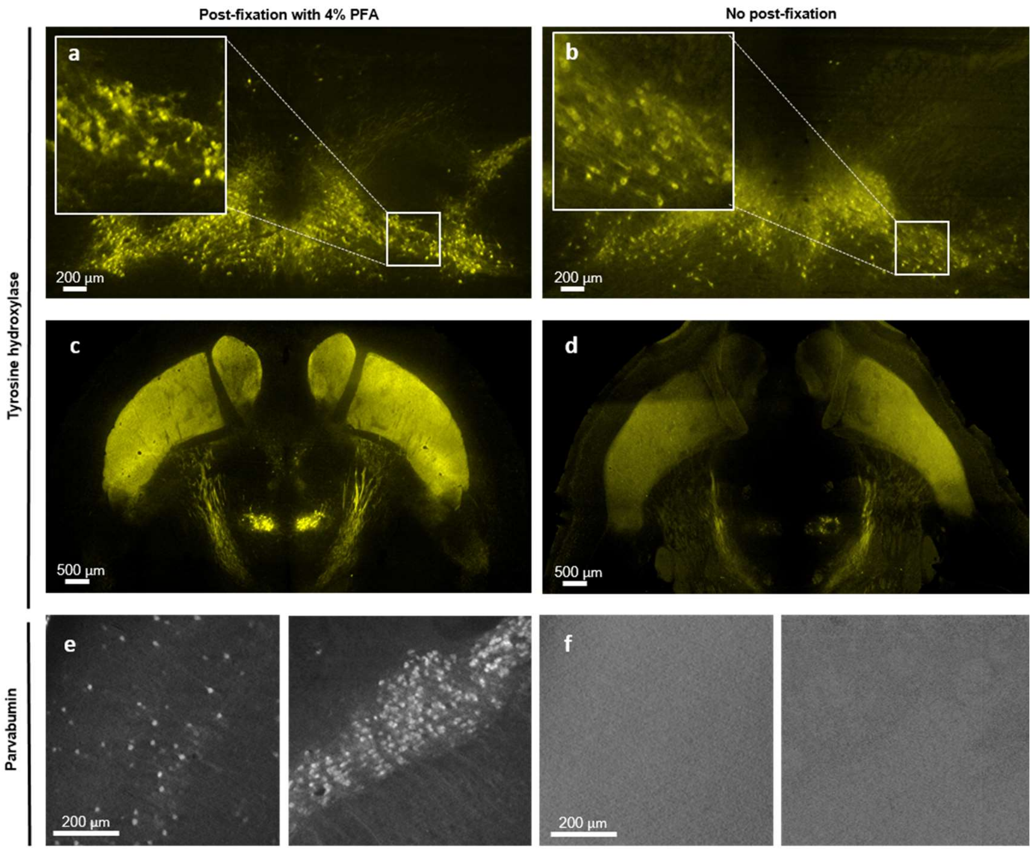


**Figure S6: Immuno-labeled whole mouse brain without post-fixation exhibits faint or negative staining.** Light-sheet images of a FFPE mouse brain stained with tyrosine hydroxylase (TH) and parvalbumin (PV) with (**a, c, e**) or without (**b, d, f**) post-immunolabeling fixation. Without post-fixation, the TH signal is fainter (**b, d**), and PV signal is completely absent (**f**).

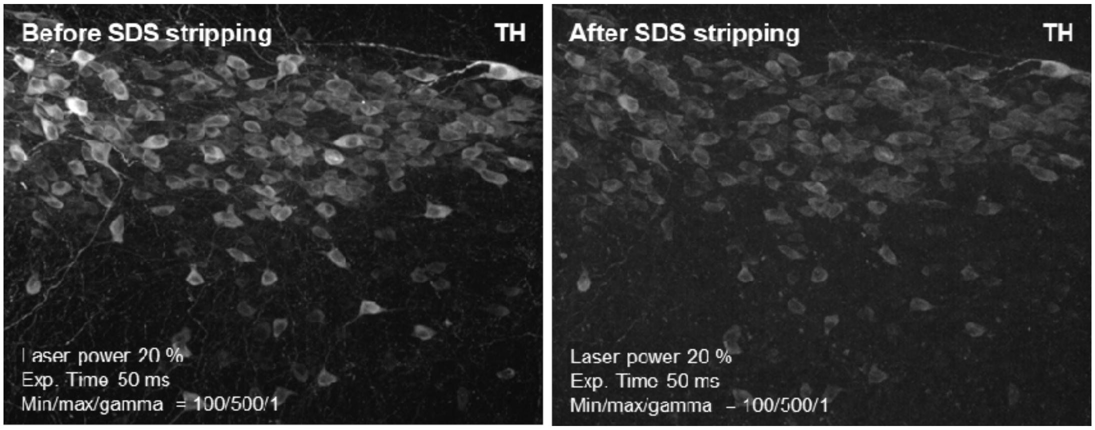


**Figure S7: Post-immunolabeling-fixed TH signal remains after SDS stripping.** A 2-mm-thick FFPE mouse brain specimen was stained with TH and then underwent post-immunolabeling fixation with 4% PFA. Multi-point confocal images were taken before and after SDS stripping. Laser power, exposure time, and display parameters are indicated.

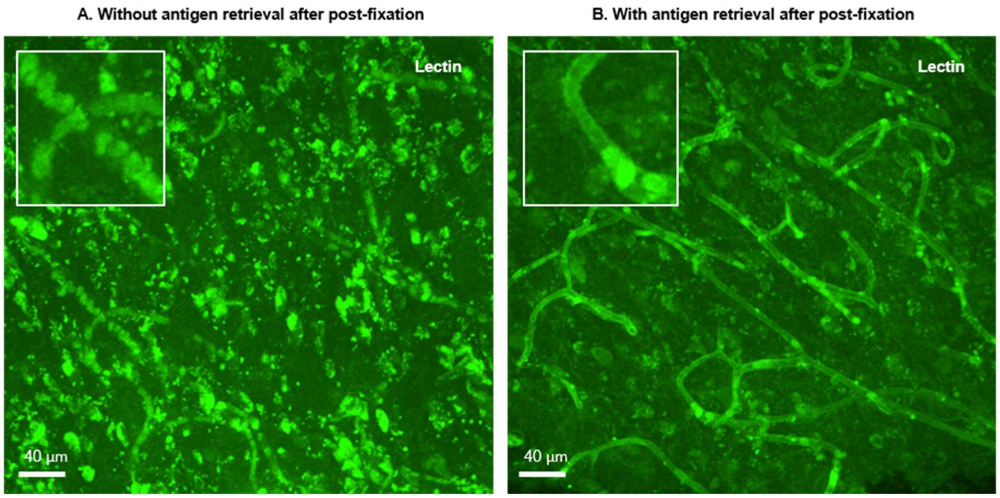


**Figure S8: Classic heat-induced epitope retrieval (HIER) facilitates relabeling after post-immunolabeling fixation.** A second round of staining, using lectin to label blood vessels, was performed on human brain tissues that had undergone an initial immunolabeling and post-fixation process. For the sample without HIER before lectin staining (**a**), only autofluorescence of blood cells is observed. In contrast, for the sample with HIER before lectin staining (**b**), the endothelium of blood vessels is clearly labeled.

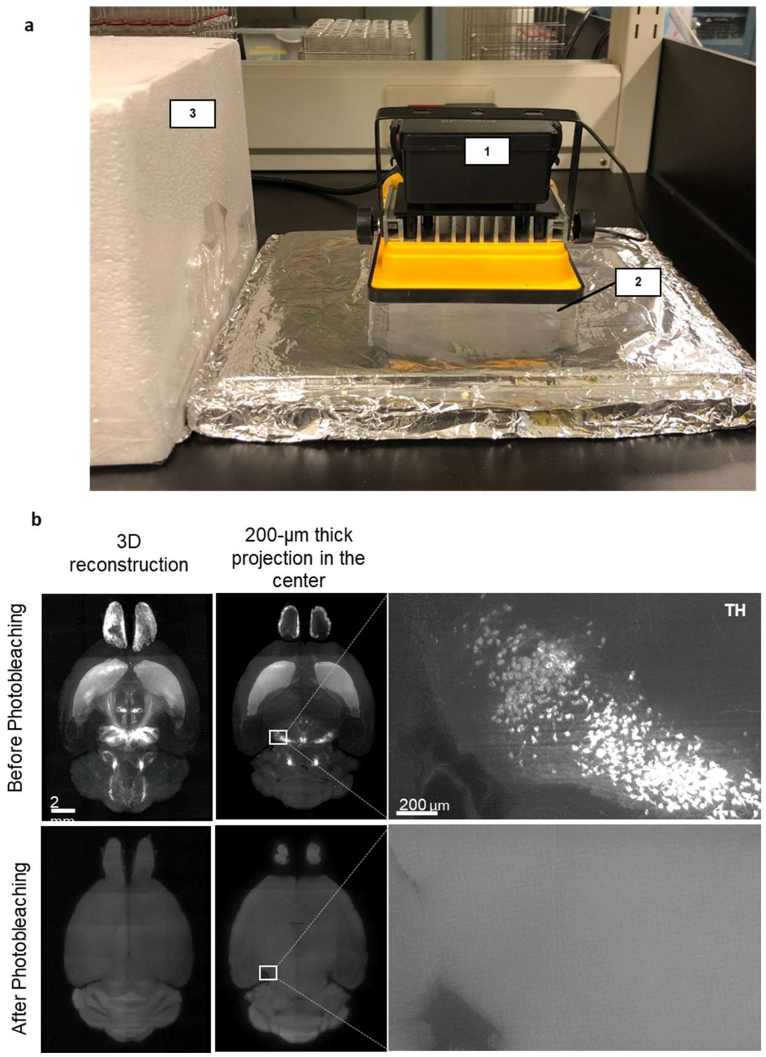


**Figure S9: Photobleaching in HIF-Clear+.**

**a.** A representative photobleaching apparatus. The 100W projection lamp with an LED array (1), the multi-well plate containing RI-matching solution and the sample (2), and the cover for the apparatus when the lamp is turned on (3).

**b,** Light-sheet images of a FFPE mouse brain stained with tyrosine hydroxylase (TH) before (upper row) or after (bottom row) white light LED photobleaching.

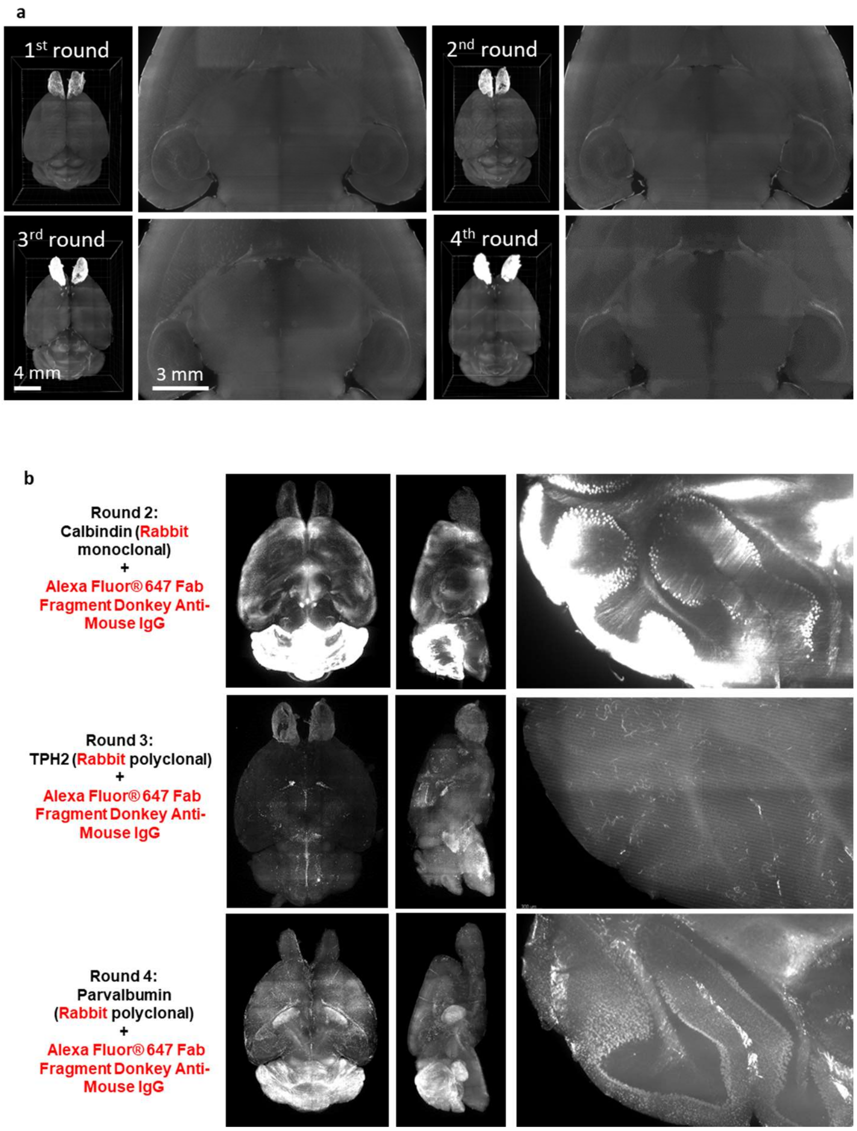


**Figure S10: Examination of issue structural integrity and secondary antibody cross-reactivity of HIF-Clear+.**

**a,** Light-sheet images of a FFPE mouse brain after each round of immunolabeling. 3D reonstruction images and magnification of the optical section in the center are shown.

**b,** The second to fourth rounds of multi-round staining shown in Figure 3. Antibodies used, Projection images of the horizontal and sagittal views, and magnification of the cerrballur region are shown. In the three sequential rounds, rabbit primary antibodies and the same secondary antibodies were utilized. Our observations under a 3.6x objective (NA = 0.2) did not reveal any colocalization with the staining from the previous round. The results suggest that cross-reactivity is not significant.

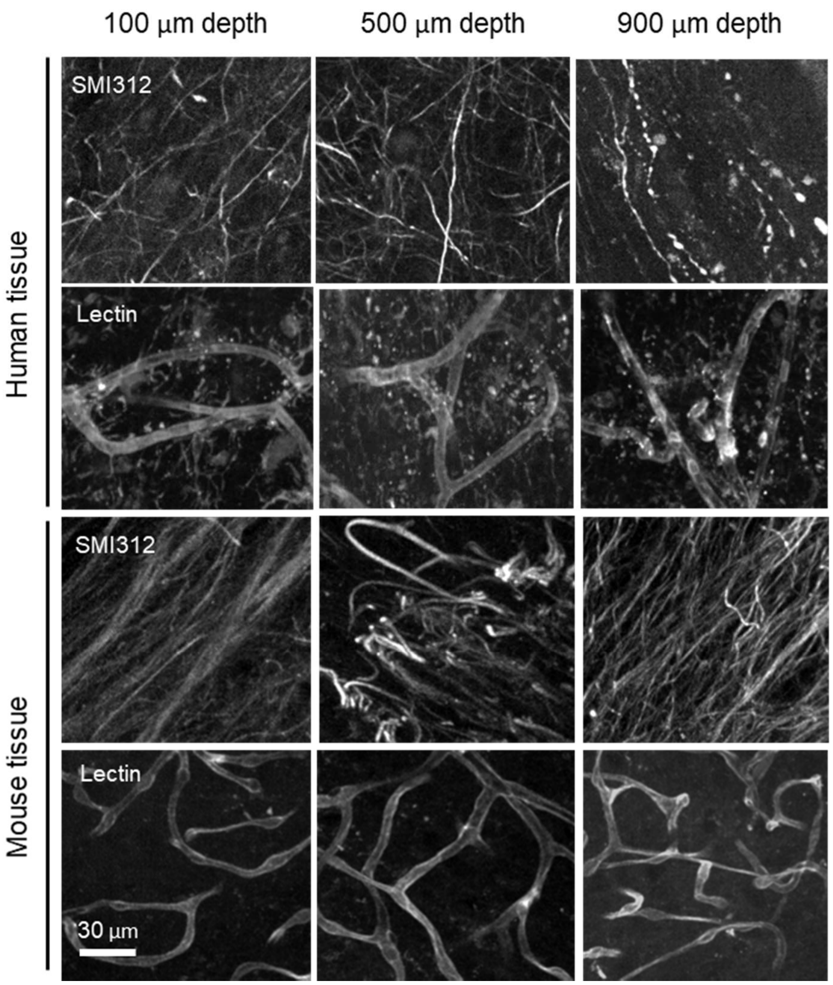


**Figure S11: Comparison of staining qualities of human and mouse brain tissues.** Multi-point confocal images of TH- and Lectin-labeling in 1mm-thick human and mouse brain tissues at depths of 100 um, 500 um, and 900 um are shown.

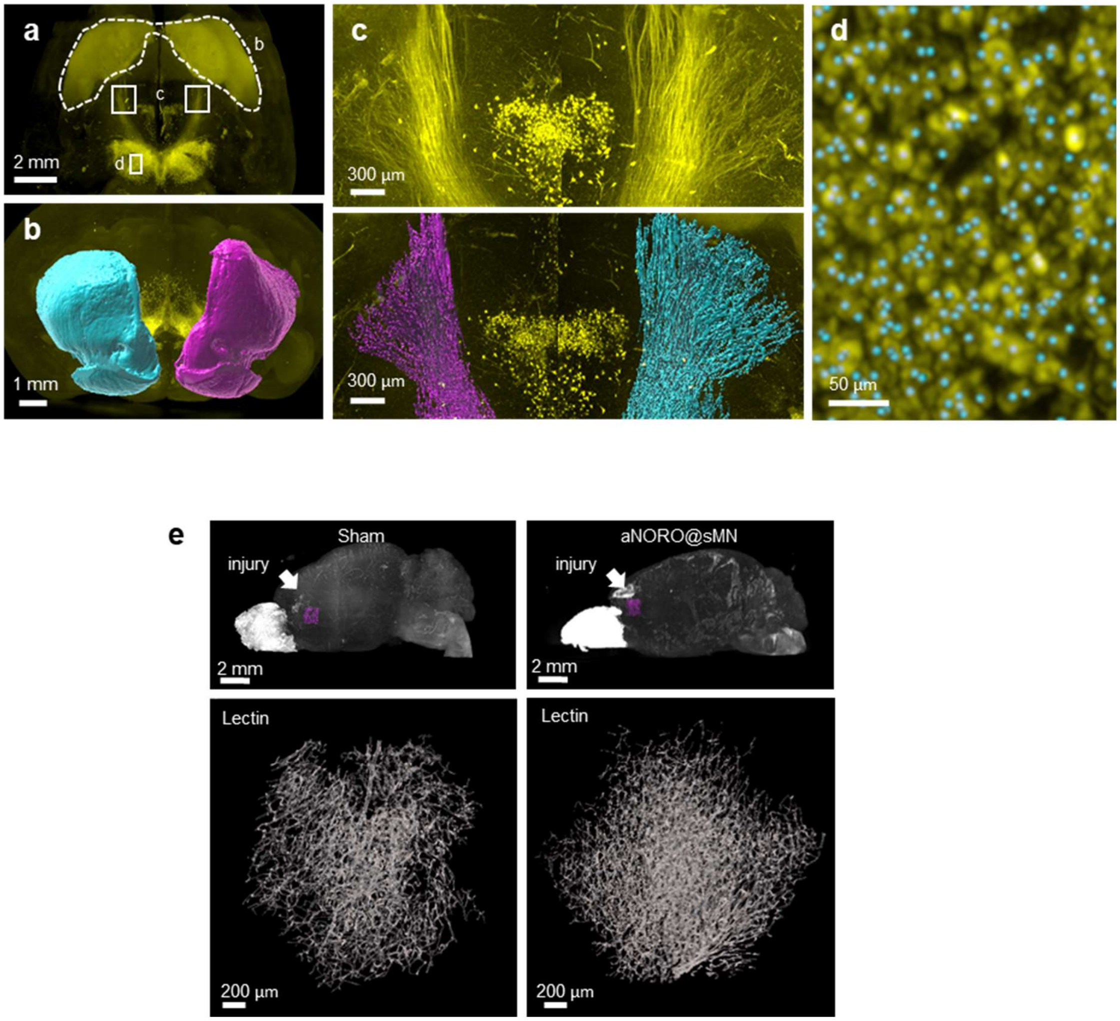


**Figure S12: Segmentation of dopaminergic regions and blood vessels in PFA-SDS TBI mouse brains.**

**a,** Whole brain projection light-sheet image focusing on TH-positive regions.

**b,** Frontal view of segmented striatum (dashed outline in **a**). Magenta: the injured side; cyan: the contralateral side.

**c,** Horizontal views of the nigrostriatal fiber tract (paired white boxes in **a**); fluorescence (top) and segmented (bottom) images are shown. Magenta: the injured side; cyan: the contralateral side.

**d,** Detection of dopaminergic cells in the substantia nigra (SN). Magnification of the area marked area in **a** (white box), including the TH fluorescence signal (yellow) and segmented cell spots (cyan dots).

**e,** Reconstruction of blood vessels in PFA-SDS TBI brains with or without treatments. The regions of interest (ROIs) selected for assessment of angiogenesis are indicated in the top row (magenta boxes, directly below the injury site). 3D reconstructions of blood vessels are shown in the bottom row.

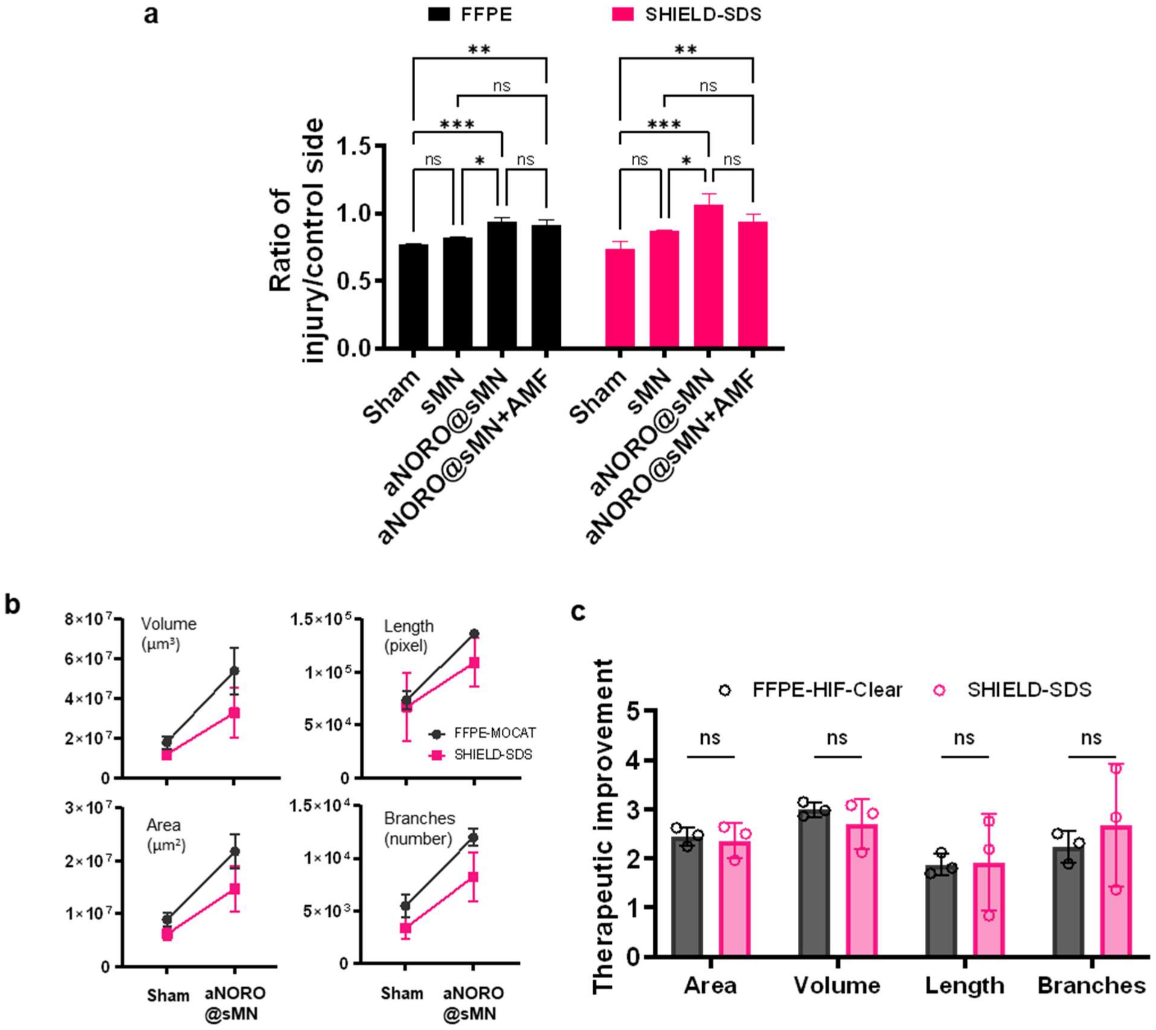


**Figure S13: Quantifications of FFPE-HIF-Clear and PFA-SDS samples result in identical statistical trends.**

**a.** Quantification of FFPE-HIF-Clear and PFA-SDS brains reveals identical inter-subgroup differences. n = 3 (three indicators: striatum volume, nigrostriatal fiber tract volume, SN cell number), mean ± SD, * $p < 0.05$, ** $p < 0.01$, *** $p < 0.001$, **** $p < 0.0001$, one-way ANOVA with Tukey's multiple comparison test.

**b,** Quantification of blood vessel volume, blood vessel surface area, number of blood vessel branches, and blood vessel length. n = 3, mean ± SD.

**c,** Comparison between the quantification results for the FFPE-HIF-Clear and PFA-SDS mouse brain samples. Multiple independent *t* tests were performed. n = 3, ns = not significant; p = 0.93, 0.87, 0.93, and 0.92 from left to right, respectively.

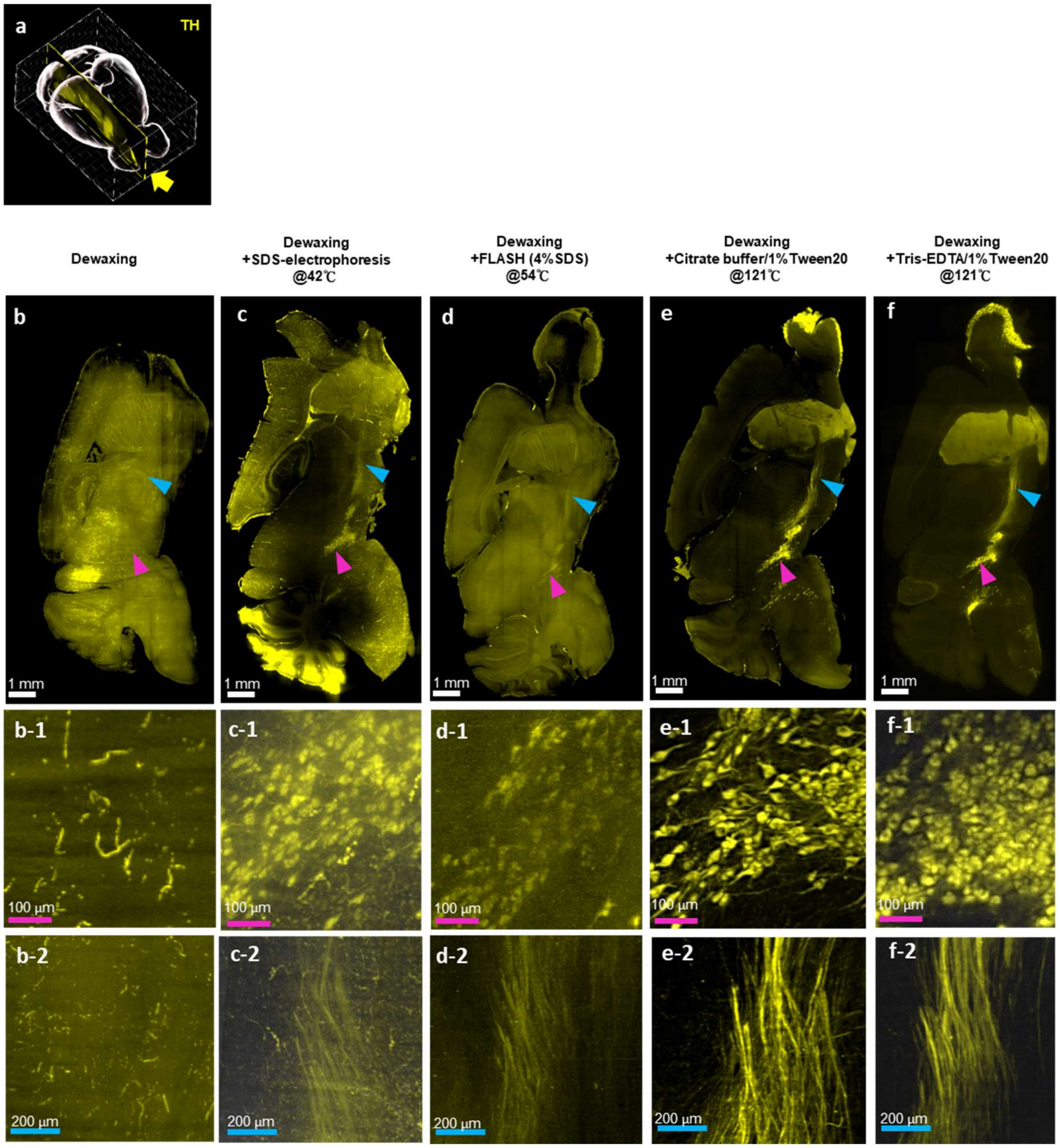


**Figure S14: Effect of various antigen retrieval conditions on FFPE mouse brains.** Light-sheet images of a FFPE mouse brain hemisphere stained with TH antibodies to evaluate the effect of different antigen retrieval methods.

**a**, The position of the Y-Z plane (yellow arrow) shown in **b, c, d, e** is indicated.

The substantia nigra (magenta arrowhead) and nigrostriatal fiber tract (cyan arrowhead) are indicated. In the mouse brain not subjected to antigen retrieval (**b**), TH signal is not detectable. The locations where TH signal should highlight the substantia nigra and nigrostriatal fiber tract are indicated by the colored arrowheads. **b**-1, **c**-1, **d**-1 and **e**-1 show magnifications of the substantia nigra indicated by the magenta arrowheads in **b**, **c**, **d** and **e**. In **b**-2, **c**-2, **d**-2 and **e**-2, magnifications of the nigrostriatal fiber tract indicated by cyan arrowheads are shown.

**Supplementary video 1:** Spatial distribution of the six biological markers shown in Fig. 4.

**Supplementary video 2:** Spatial relationships of the tumor and surrounding astrocytes in Fig. 7A to E. The segmented astrocytes are colored according to the cell-to-tumor distance code.

**Supplementary video 3:** 3D visualization of the segmented blood vessels shown in Fig. 7L.
